# Online Bayesian Analysis with BEAST 2

**DOI:** 10.1101/2022.05.03.490538

**Authors:** Remco Bouckaert, Lena Collienne, Alex Gavryushkin

## Abstract

There are a growing number of areas, e.g. epidemiology and within-organism cancer evolution, where re-analysing all available data from scratch every time new data becomes available or old data is refined is no longer feasible. All these and related areas can benefit from online phylogenetic inference that can booster previous data analyses.

Here, we make the case that adding/removing taxa from an analysis can have substantial non-local impact on the tree that is inferred, both in a model based setting, as well as for distance based methods. Consequently, online phylogenetic algorithms may require global updates of the trees and other parameters, a task that in general is highly non-trivial.

Motivated by this observation, we designed an online algorithm that benefits from a parallelism in a Bayesian setting that is substantially more efficient than re-running the analysis from scratch. Furthermore, our algorithm is not sensitive to the number of sequences added, allowing the sequence data to grow/be refined iteratively. We show how this approach can be used in a maximum likelihood setting, and – apart from adding/removing new sequences – demonstrate a number of practical alternative use cases of our algorithm, including how to break up a single (offline) large analysis to get results faster.

An open source implementation is available under GPL3 license as the ‘online’ package for BEAST 2 at https://github.com/rbouckaert/online and a tutorial at https://github.com/rbouckaert/online-tutorial.

## 1. Introduction

Rapid development of molecular sequencing technologies in recent years highlights the importance of scalable computational approaches to data analysis. *Online* (also known as dynamic) algorithms are algorithms that are able to update the analysis results when data is updated. Ultimately, online algorithms enable one unified approach to data processing and analysis continuously in parallel to experiment design and data collection. Typical examples of online data updates include data point additions, removal, and modifications. Although the question of when an (*offline*, also known as static) algorithm has an efficient online version received considerable attention in the theoretical computer science literature (Karp, 1992; Albers, 2003; Borodin and El-Yaniv, 2005), the problem still remains largely unsolved. Indeed, the majority of results in this area constitute examples of online algorithms (Christensen et al., 2017; Mairal et al., 2010; Psaraftis et al., 2016) while a general technique applicable to a wide range of (offline) algorithms is still lacking. So designing new online algorithms for specific problems is a highly non-trivial task (Gavryushkin et al., 2015, 2016).

Online algorithms have received a fresh wave of attention in recent computational biology literature including phylogenetics (Gill et al., 2020; Dinh et al., 2018; Fourment et al., 2018), where effectively no practical online inference method is available despite the need for such methods in applications including epidemiology and cancer evolution. For example, the tsunami of sequencing data of the SARS-CoV-2 virus over the last two years has demonstrated how much more empowered our computational approaches could have been were an online phylodynamics platform available capable of extending results of analyses with new sequences arriving in batches from the outbreak’s hotspots around the globe (Turakhia et al., 2021). Without such a platform, even the virus sequences isolated from COVID-19 patients in Australian and Aotearoa New Zealand were too many to analyse in a full Bayesian framework and subsampling was required (Douglas et al., 2020, 2021a). In single-cell cancer phylogenetic, the ever growing data sizes (in terms of both sequence length and numbers of sequenced cells) without established data processing technologies (Moravec et al., 2021) implies that the same analyses need to be rerun many times for slightly changed data, e.g. with different data filtering thresholds (Moravec et al., 2021). An online approach, if it was available, in this context would bring a significant reduction in computational costs. In fact, implementing Bayesian phylogenetic inference methods in the online framework to include a wide range of evolutionary models has been identified as in important future directions (Fourment et al., 2018).

Bayesian phylogenetics with its heavy reliance on Markov chain Monte Carlo (MCMC) algorithms is an area where online approaches should be feasible to design and implement, because MCMC chains can be continued to accommodate changes in data. Indeed, this approach has been shown to be fruitful (Gill et al., 2020). However, the resumed chain still takes significant time to converge, leaving the question of whether the potential of MCMC algorithms can be further exploited to significantly increase the efficiency of online Bayesian tree inference methods. Another promising approach to update Bayesian posterior samples in the online framework is sequential Monte Carlo (SMC) (Dinh et al., 2018). Although the current implementation suffers from scalability issues allowing only a limited number of sequence additions, the potential of online SMC methods is clear (Dinh et al., 2018).

In this paper, we present a full Bayesian approach to phylogenetic inference in the online framework where online modifications are (existing) sequence removals and (new) sequence additions. That is, our method is capable of updating a Bayesian posterior tree (and other parameter) sample after a set of existing sequences (taxa) is removed and/or a set of new sequences (taxa) is added. The update operation requires significantly less time to be performed than running the analysis from scratch. Furthermore, the update operation outputs a converged chain (assuming a converged chain was available before sequence removals/additions), hence the number of sequence removals/additions is unlimited.

Briefly, our update operation is performed in three steps: (1) remove obsolete sequence and place new sequences on the existing trees; (2) perform local MCMC update of the part of the trees where the new sequence are placed; (3) perform global MCMC to accommodate any necessary non-local changes to the trees (and other parameters). Our online algorithm is implemented in the BEAST 2 (Bouckaert et al., 2019) framework, so it comes with a wide range of evolutionary models, limited only to those available in BEAST 2. As a by-product of the work presented in this paper, we discuss how our method implies an efficient online phylogenetic inference algorithm in the maximum likelihood framework.

Finally, we study the question of online algorithms for non model-based phylogenetic reconstruction methods. Specifically, we consider the neighbour joining (NJ) and the unweighted pair group method with arithmetic means (UPGMA) tree reconstruction algorithms, and discuss the question of when a distance-based method has an efficient online version. Through so-called stability analysis we demonstrate that both NJ and UPGMA algorithms are not ‘local’ meaning that online versions for them are hard to design.

## 2. Two motivating examples

The aim of efficiently updating a Bayesian phylogenetic posterior sample following taxa removals/additions would have been relatively straightforward to achieve if the posterior of a full set of taxa *X* marginalised to take out some subset of taxa *S* would be equal to the posterior under the same model for only the set of taxa *X\S*. However, we will argue in this section that this is not true. Specifically, we will provide two examples where the two distributions are different. This fact will guide the design of our online algorithm in the sections that follow.

### 2.1. The Chicken problem

Our first example of this phenomenon is summarised in Fig. 1, which shows the tree distributions visualised using DensiTree (Bouckaert and Heled, 2014) for a dataset with 10 taxa^1^. The first panel shows the tree distribution sample obtained from the full set of 10 taxa. The second shows the analysis with 9 taxa when the human sequence is removed. Noticeable is the placement of the chicken taxon in the tree. While in the 10 taxon analysis the chicken is monophyletic with frog, loach, and carp, the 9 taxon analysis is not so certain, and makes the clade monophyletic with only about 50% posterior support. The other 50% support goes to placing chicken with the mammals, or as an out-group of the other 9 taxa.

**Figure 1.**
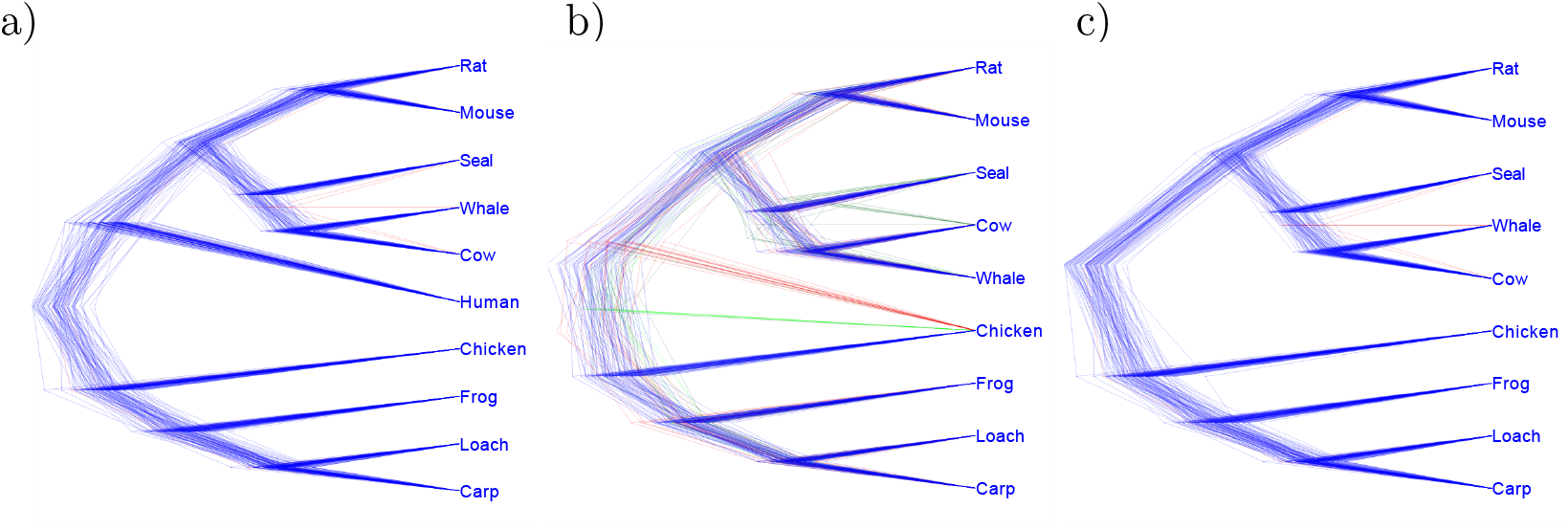
Illustration of the chicken problem: a) full distribution, b) tree distribution when human sequence is removed, c) the marginal distribution when the human sequence is removed.

The third panel shows the 10 taxon tree distribution with the human branch filtered out, that is, the 10 taxon tree posterior with humans marginalised out. This tree distribution differs from the 9 taxon analysis distribution, since the clade with chicken, frog, loach, and carp is 100% monophyletic.

### 2.2. The Bear-Cat-Dog (BCD) problem

Our second example is summarised in Fig. 2. We consider the following three taxa from a mitochondrial alignment of 33 mammals (Yang and Yoder, 2003): bear *Ursus americanis* (B), cat *Felis catus* (C), dog *Canis familiaris* (D), and horse *Equus caballus* (H). It turns out that the distance between B and C is much smaller than that of D and B as well as D and C. Any reasonable phylogenetic method therefore will reconstruct the tree as ((B,C),D). Here, we use the analysis used in (Bouckaert et al., 2019) with all sequences but B, C, and D removed. It uses a Yule tree prior, strict clock model, and HKY substitution model with gamma rate heterogeneity with 4 categories. The result is that ((B,C),D) has 54.5% posterior support, while ((B,D),C) has 34.4% and ((C,D),B) has the remaining 15.1% support (Fig. 2a).

**Figure 2.**
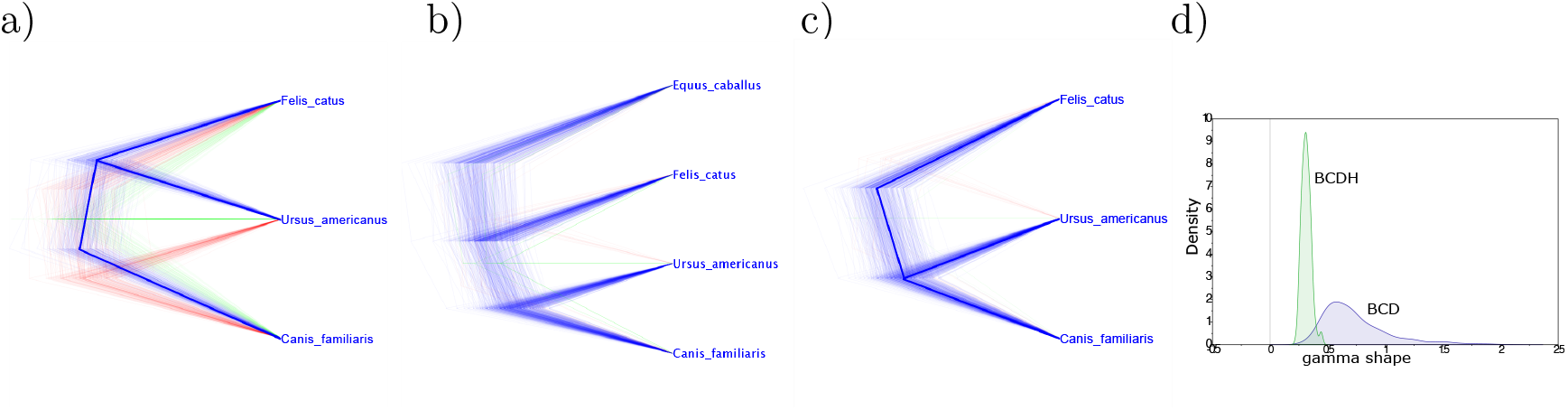
Illustration of the BCD problem: a) distribution over B = bear, C = cat and D = dog, b) tree distribution when horse sequence (H) is added, c) the marginal distribution over B, C, and D when H is removed, d) marginal distribution of gamma shape parameter.

When we add a sequence of horse (H) to the data, the subtree ((B,D),C) gets 100% support, so the marginal distribution of the analysis over the four taxa differs substantially from the analysis of the three taxa B, C, and D alone (Fig. 2). The addition of the H sequence leads to a slight change in the estimate of the kappa parameter of the HKY model, while the birth rate remains virtually unaltered. However, this addition considerably changes the estimate of the shape parameter for gamma rate heterogeneity (Fig. 2d), which is reduced from an average of 0.7391 when including only B, C, and D to 0.3139 when including H as well.

These two examples allow us to conclude that the addition of just one sequence can have a significant effect on model parameter estimates, including the tree estimate. Specifically, the knowledge about a posterior distribution over a tree-space with *k* taxa is not sufficient to place an additional taxon unless it is allowed to alter the *k*-taxa trees. The chicken example illustrates uncertainty around placing the chicken taxon with respect to the root. In general, this uncertainty is present at internal nodes as well, and by adding taxa the uncertainty can be reduced in a similar fashion as how the chicken placement is resolved once the human sequence is added.

One could argue that these changes in topology are still localised, but the BCD example implies that online phylogenetic inference algorithms have to also account for site model parameter changes. These site model parameters can influence the whole of the phylogeny, so changing them in the online framework can lead to internal node height changes and even tree topology changes as in the BCD example.

## 3. Algorithm

With the examples from the previous section in mind, we are now ready to describe our online algorithm that efficiently updates a given Bayesian posterior sample when new taxa (sequences) are added and/or existing taxa are removed. Motivated by numerous applications, e.g. the ongoing COVID-19 outbreak studies in epidemiology, our algorithm is designed to accommodate additions/removals of multiple taxa (rather than one by one).

We assume that we have a phylogenetic analysis (using e.g. BEAST 2) for a certain number of sequences already completed, and that we have a trace of the MCMC states. We ignore burn-in, and only consider the MCMC states from the stationary part of the MCMC chain. Our goal is to run a new MCMC analysis with extra taxa in the alignment and/or some of the existing taxa removed. To be able to remove existing taxa in the online framework is important because in practice misaligned and error ridden sequences may only be identified after an analysis is performed.

Building on the approach of Gill et al. (2020), which takes the state at the end of an MCMC run, we instead consider a sufficiently large number *n* of stored MCMC states evenly spaced among the stored MCMC states from the stationary part of the chain (more about the size of *n* below). For each such state, we create a new sample starting at the stored state, resulting in an independent sample from the new posterior of interest. Since these *n* instances can be run independently, this is implemented in the obvious (embarrassingly) parallel way.

The creation of an independent sample from the larger space has three distinct phases (Fig.3), which we describe in detail below: 1) Placing (and/or removing) taxa in the tree, 2) Performing local MCMC, and 3) Performing full MCMC.

**Figure 3.**
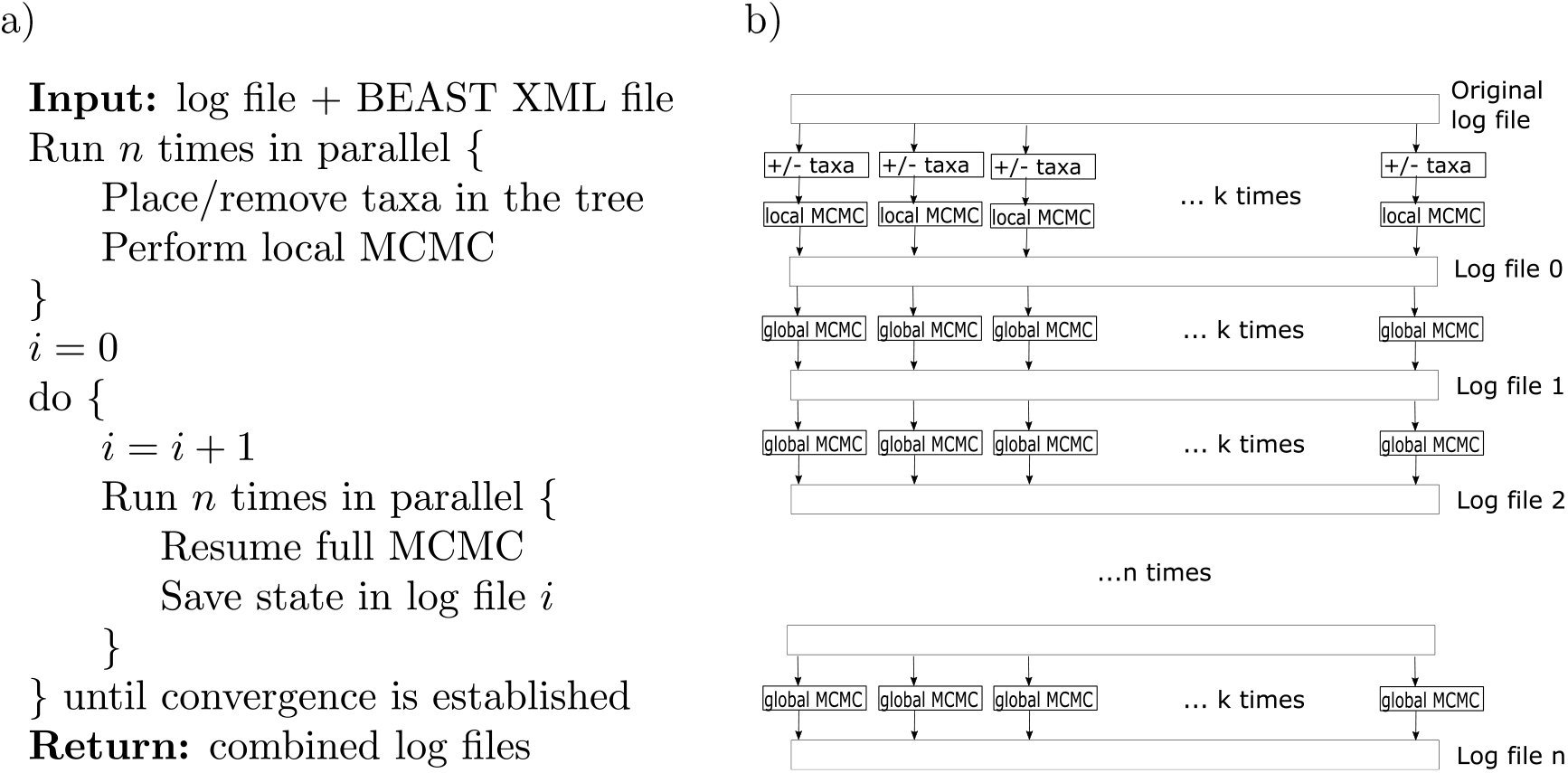
Algorithm 1: Online Bayesian inference algorithm. a) pseudo code, b) schematic representation of processes and files involved.

As a result, given sufficiently many CPU cores, the *n* samples can be generated in a much shorter time frame than a full MCMC run can. We will run the full MCMC chains at least as long so to generate independent samples to be able to establish convergence. This gives us at least 2*n* independent samples. If we aim for an effective sample size (ESS) of *E* = 200, then *n* = *E/*2 = 100 states is sufficient, so this is the starting point of our algorithm (and this number remains fixed for all online iterations of the algorithm). By combining two samples at sufficiently large intervals apart, we double the ESS and get the desired ESS of *E*. Alternatively, when the number of available cores is insufficient to run *E/*2 threads in parallel, a lower number for *n* combined with a higher number of samples can be used to obtain the desired ESS.

### 3.1. Placing new taxa in the tree

Adding taxa to existing trees in a Bayesian setting can be done efficiently with PPlacer (Matsen et al., 2010). Here, we take a slightly different approach by doing a binary search through the tree for each of the taxa to add such that it maximises the posterior probability. The result will be a tree with all new taxa added and fairly high posterior support.

To add a single taxon, we start by adding a new root node whose children are the new taxon and the old root. We denote thus obtained tree by *T*_*root*_. The height of the new root node is set to the old root height times (*m* + 1)*/m* where *m* is the number of nodes in the old tree. We then calculate the posterior *p*_*root*_ for this tree. Next, we move the new root to half way the left branch of the old root giving tree *T*_*left*_ and calculate posterior *p*_*left*_, and likewise for the tree *T*_*right*_ with the new taxon placed along the right branch, giving *p*_*right*_. If *p*_*root*_ *> max*(*p*_*left*_, *p*_*right*_), we stop and return *T*_*root*_. If *p*_*left*_ is the highest posterior, we iterate this process with *T*_*left*_ acting as *T*_*root*_, otherwise we iterate with *T*_*right*_. We keep on descending in this way along the branches down the tree, until both left and right posteriors associated with the proposed placement are lower than the running posterior, or a branch leading into a tip is reached. This process is repeated until all taxa have been added to the tree.

Since posterior calculation only involves recalculating tree likelihoods, which can be done efficiently by caching previous partial posterior calculations, only the part of the tree from the running placement branch to the root needs recalculation using Felsenstein’s peeling algorithm. In the worst case this taxon placement algorithm is quadratic in the number of tree branches (when it requires placement of a taxon in the cherry clade of a caterpillar tree), but in practice the algorithm is efficient enough that run time is negligible compared to performing MCMC in the two following stages.

Note that with the placement of each taxon, potentially other parameters change as well. The dimension of each such parameter (e.g. substitution rates along the branches) needs to be increased by 2 after every taxon placement, since each additional taxon results in two extra branches.

- For substitution rates, a branch that is split in two due to addition of an internal node receives the same rates (i.e. the original branch rate is duplicated) and the rate for the branch to the taxon is set to 1.
- For boolean parameters (e.g. the rate change indicator for random local clocks), all newly added entries are set to false.
- For integer parameters (e.g. rate categories for uncorrelated relaxed clock (Drummond et al., 2006)) new entries are initialised in the middle of their range.

After all new taxa are added and, if necessary, some of the existing taxa are removed, we run the following two steps in parallel for each of the *n* trees.

### 3.2. Performing local MCMC

After placing taxa as described above, it is necessary to rearrange the tree due to unforeseen alterations similar to the chicken and BCD-problems (Section 2). However, since the biggest changes can be expected around the area where the new taxa were placed, initially, we only allow proposals that perform updates of the tree in the neighbourhood of the inserted internal nodes.

The neighbourhood is defined as the new internal node’s two children, its parent, and the parent’s children (which include the new node). This is the neighbourhood used in our computational experiments and the algorithm’s performance evaluation. Not only the tree, but parameters associated with branches, such as, for example, clock rates in the relaxed clock model (Douglas et al., 2021b) or random local clock (Drummond and Suchard, 2010) associated with branches in the neighbourhood are allowed to be updated.

### 3.3. Performing full MCMC

The two examples above (the BCD and Chicken problems) suggest that this local MCMC step might not be enough to provide an unbiased sample from the posterior distribution over bigger trees because e.g. global parameters need to be update and they in return can impact the trees. Indeed, at the end of the local MCMC chain we finish in a state that has a higher posterior than at the start, but other parameters in the model, as well as other parts of the tree, interact in such a way that the sample is biased. To get an unbiased sample, it is necessary to sample the full state of the model, including all site model parameters, other clock model parameters, other parts of the tree and tree prior parameters. So, we run a full MCMC chain *n* times in parallel for each of the local MCMC samples, and combine the logs as final result of one iteration of our online algorithm.

The local MCMC step is important for overall performance of the online algorithm in terms of *efficiency*. Indeed, the MCMC proposals of the local step of the algorithm require on average significantly less computation than the full MCMC proposals. This is because posterior calculations are dominated by tree likelihood calculations and for local tree proposals these can be done efficiently by caching and recalculating only for the part of the tree that changes plus the nodes leading from there up to the root. In contrast, in the full MCMC step, every update of site model parameters or mean clock rates requires recalculation for all nodes in the tree, which is more expensive.

However, the full MCMC step is crucial to *guarantee convergence*, i.e. to guarantee that the final sample is a sample from the true posterior distribution. Indeed, the fact that plain (offline) MCMC converges to the true posterior distribution implies the same is true for our online algorithm as running multiple MCMC chains simply provides multiple individual samples from the same posterior distribution. Computationally this is ensured by running the full MCMC step for a sufficiently long time, as we discuss in the next section.

### 3.4. Stopping criterion

Ideally, an online algorithm full Bayesian phylogenetic inference algorithm should terminate automatically before taking in new sequences and going to the next online iteration. In this section we describe how this process is implemented in our algorithm, however, this task is essentially equivalent to the notoriously hard problem of deciding whether an MCMC algorithm has converged (Cowles and Carlin, 1996; Vehtari et al., 2021). We do not suggest that we provide an ultimate solution to this stopping problem, but that our online algorithm can benefit from general research in this area.

There are a number of statistics that can be calculated to compare a sample at the end of the full MCMC chain consisting of the states of all *n* MCMC instances, and a states that are saved periodically during the full MCMC chain. The literature offers the Gelman-Rubin statistic (Gelman et al., 1992; Brooks and Gelman, 1998), the split-R variant (Vehtari et al., 2021), and others (Vats and Knudson, 2018; Cowles and Carlin, 1996). Our implementation allows automatic stopping based on the statistics mentioned above, as well as a fixed limit on the number of cycles in step 3 of the algorithm, which then allows comparison of samples at the last cycle with samples at earlier samples. If these samples are sufficiently close based on the estimate of the mean, Kolmogorov-Smirnov statistic (Massey Jr, 1951), cumulative difference of kernel density diagrams, and have little correlation, it is reasonable to assume, and so do we in our algorithm, that the MCMC has converged. However, if many parameters are involved, these statistics may sometimes fail to reliably diagnose convergence. Alternatively, manual inspection of visualisation of these samples in Tracer can give an insight into how much samples differ.

Unfortunately, we did not find a satisfactory criterion to fully automatically decide when to stop the *n* full MCMC chains: the statistics seemed either too liberal like the Gelman-Rubin statistic, while others were overly conservative like the KS statistic. This is not surprising, since it is generally necessary to use multiple criteria to judge convergence of MCMC chains (Cowles and Carlin, 1996; Vehtari et al., 2021). It also implies that ultimately human judgement still is required to reliably determine convergence.

## 4. Validation

We performed a well calibrated simulation study to validate the correctness of our implementation. As a practical validation, we perform an online analysis of SARS-CoV-2 sequences from community outbreaks in New Zealand.

### 4.1. Implementation correctness

We performed a well calibrated simulation study sampling 100 tip dates randomly from the interval 0 to 1. The taxa are ordered by tip date to allow us to remove the *k* youngest tips, and then add them back in an online fashion.

We use a coalescent tree prior with constant population size log-normal(*µ* = 1, *σ* = 1.25) distributed, an HKY model with kappa log-normal(*µ* = 1, *σ* = 1.25) distributed, and gamma rate heterogeneity with four categories with shape parameter exponentially distributed with mean=1 and frequencies Dirichlet(1,1,1,1) distributed. Further, gamma is lower bounded by 0.1 (Bouckaert, 2020) and frequencies are lower bounded by 0.2 to prevent atypical parameter values. We use a strict clock where the clock rate times tree height has a tight normal(*µ* = 1, *σ* = 0.05) prior. Sampling 100 instances from this distribution using MCMC in BEAST (Bouckaert et al., 2019), we get a range of tree heights from 1.03 to 8.8 with mean 1.6 (note that due to the tips being sampled from 0 to 1, the tree height is lower bounded by 1) and we get a range for the clock rate of 0.1 to a fraction over 1 in our study. With these trees, we sample sequences of 1000 sites using the sequence generator in BEAST 2.

From these 100 taxon samples, we removed the newest *k* sequences with *k* = 10, 20, 30, 40, 50, run a full MCMC with 20 million samples, which is sufficiently long to get effective sample sizes of around 200 for the posterior. Then, we ran our online algorithm with the full 100 taxa for 50 thousand samples for each of the 200 states obtained from an MCMC run, running 500 MCMC analyses (100 each for 50, 60, 70, 80, and 90 taxa) and 500 online analyses (each for 100 taxa) in total.

Table 1 shows the coverage of true parameter values (and some other statistics) used to simulate the sequence data by the 95% highest probability density (HPD) intervals estimated after running the online algorithm. With 100 experiments, the 95% HPD of the binomial distribution with p=0.95 ranges from 91 to 99 inclusive. Only the population size estimate when 10 taxa are added has a coverage of 100, indicating a slight over-estimate in the size of HPDs. A single such outlier can be expected when comparing this many (5×11=55) statistics, so this gives confidence the implementation is not incorrect.

**Table 1.** Coverage of the true value by 95% HPD estimates from 100 independent runs of the online algorithm. Columns indicate the number of taxa added to make a total of 100 taxa. All coverage is in the expected 91 to 99 range.

| Parameter | Taxa added |  |  |  |  |
| --- | --- | --- | --- | --- | --- |
|  | 10 | 20 | 30 | 40 | 50 |
| tree height | 94 | 94 | 95 | 91 | 93 |
| tree length | 96 | 99 | 95 | 97 | 94 |
| kappa | 95 | 96 | 99 | 96 | 98 |
| gamma shape | 93 | 93 | 95 | 96 | 97 |
| population size | 100 | 99 | 97 | 99 | 98 |
| clock rate | 96 | 92 | 95 | 93 | 93 |
| coalescent prior | 99 | 96 | 95 | 96 | 96 |
| frequencies A | 98 | 98 | 96 | 97 | 97 |
| frequencies C | 92 | 93 | 95 | 94 | 97 |
| frequencies G | 97 | 97 | 97 | 99 | 98 |
| frequencies T | 94 | 94 | 95 | 97 | 98 |

### 4.2. SARS-CoV-2 in New Zealand

We used 730 full genome sequences of the SARS-CoV-2 virus from Douglas et al. (2021a) containing samples from 11 community outbreaks in New Zealand. Genomic sites were partitioned into three codon positions, plus a non-coding region, as described in Douglas et al. (2020). For each partition we model evolution with an HKY substitution model with log-normal(*µ* = 1, *σ* = 1.25) prior on kappa, frequencies estimated with Dirichlet(1,1,1,1) prior, and relative substitution rates with Dirichlet(1,1,1,1). We use a strict clock model with log-normal(*µ* = − 7, *σ* = 1.25) prior on the mean clock rate, and as a tree prior we use a Bayesian skyline model (Drummond et al., 2005) with a Markov chain distribution on population sizes and log-normal(*µ* = 0, *σ* = 2) on the first population size.

We start with a full MCMC run with 320 taxa sampled until the end of August 2020. Then, we run our online algorithm at one month intervals until the end of December 2020. Fig. 4 shows how the tree distribution evolves during the various iterations of the last run. After the initial run, there is considerable deviation from the full MCMC run for December: node height estimates are all higher (blue crosses) and uncertainty intervals on clade heights are much larger than desired (histogram at the top of panel). Also, many monophyletic clades inferred by the full MCMC are not highly supported at the first iteration (red dots clustering at the right hand side of the panel). After the second iteration, the difference between the obtained sample and the full MCMC sample already diminishes considerably, and from 7 iterations onwards there is no further reduction in the difference between samples obtained by the online algorithm and the full MCMC sample. A full MCMC analysis takes just over a day to converge, while the online algorithm takes around 4 hours on *n* = 10 high-performance threads.

**Figure 4.**
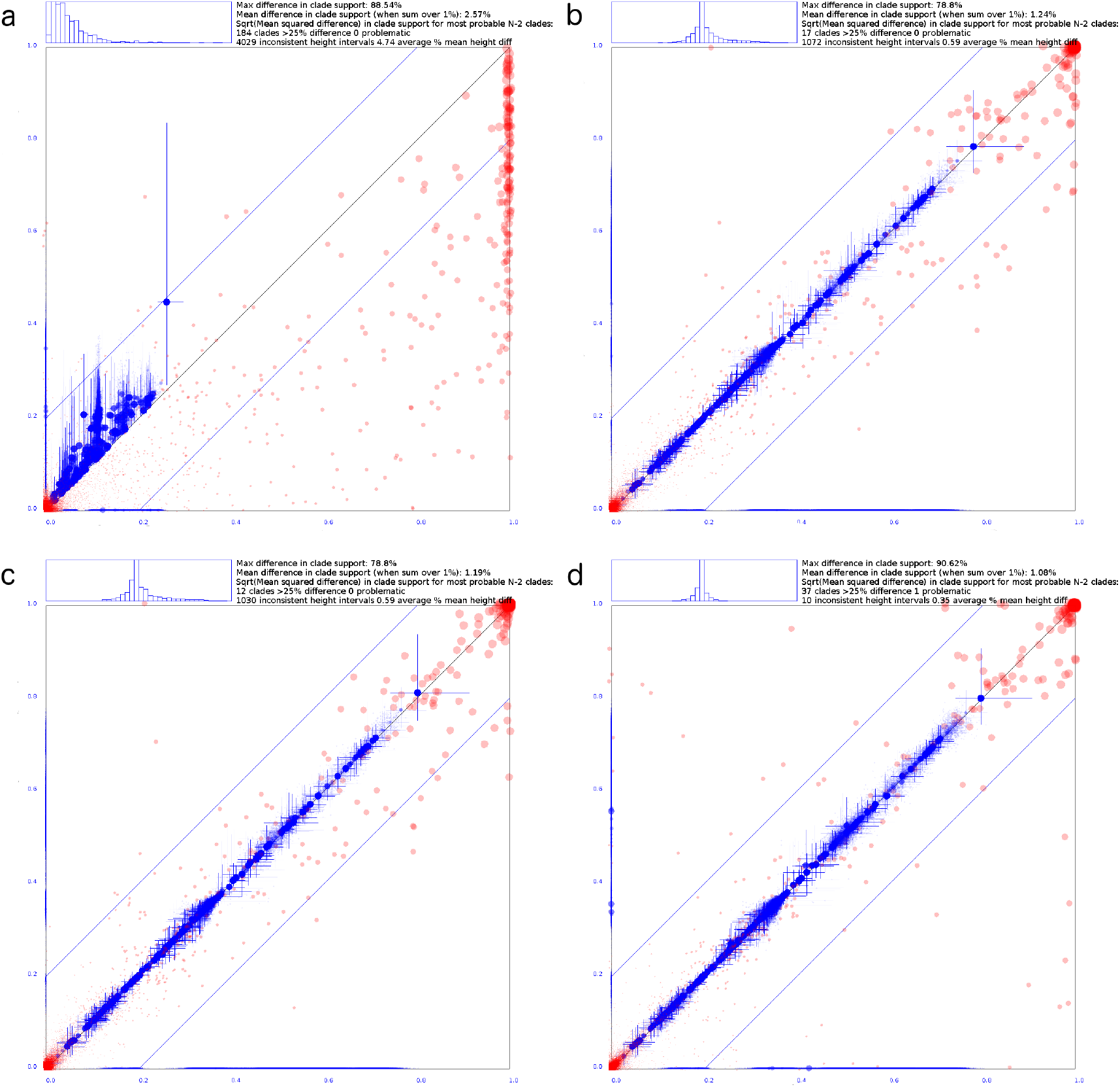
Clade set comparison of tree posterior obtained with full MCMC (horizontal axis) and after a) 1, b) 8 and c) 11 online iterations (in reading order) on the vertical axis. See Fig.S4 for intermediate cycles. The last panel (panel d) compares two independent MCMC runs for reference. Red dots indicate clade support between 0 and 1 on both axes, blue dots indicate mean clade heights with cross hairs showing the 95% HPD intervals of height estimates. The axes are scaled between zero and the highest tree height found in either tree set. Histogram at the top of each panel shows the distribution of differences in branch lengths above clades, with equal lengths in the middle, twice the length at the right hand side of the histogram, and half the length at the left hand side. After 7 online iterations the tree distribution obtained by our online algorithm becomes hard to distinguish from the full (offline) MCMC algorithm.

Some site model parameters changed significantly from the first MCMC run with the data until the end of August, and the analysis including data until the end of December. For example, the kappa parameter for the non-coding region changed from a mean of 2.47 ((1.25, 3.96) 95% HPD) to a mean of 1.84 ((1.23, 2.44) 95% HPD). The relative substitution rate per site for the non-coding region changed from a mean of 2.53 ((1.69, 3.69) 95% HPD) to 3.247 ((2.73, 3.71) 95% HPD). This yet again demonstrates that adding sequences to an analysis can have significant impact on estimates of site model parameters, and the distributions obtained from a subset of sequences can only be taken as starting point in estimating parameters for the full set, requiring global updates.

## 5. Discussion

This paper is a contribution to the active and important area of online phylogenomics, which can generally be defined as design and analysis of phylogenomic inference algorithms with data undergoing serial updates such as removals, additions, or modifications. Although such online approaches to phylogenomic inference depend heavily on the range of possible update operations, a typical scenario in practice is new sequences arriving through time, possibly in batches. One of the most recent such examples of new sequencing data arriving at scale is the epidemiological analysis of the SARS-CoV-2 outbreak, when the demand for and massive lack of online approaches in computational biology generally and in computational phylogenomics specifically has become apparent. For computation-heavy and information-rich approaches such as the full Bayesian analysis, this is felt the strongest when previous analyses were unable to converge before new sequence data has become available, and sub-optimal approaches, such as data subsampling, were used widely.

Our approach offered in this paper and motivated by this scenario is a full Bayesian phylogenetic inference method (implemented in BEAST 2), capable of incorporating new sequencing data “on the fly”. Our method passes the “minimum efficiency threshold”, that is, it is significantly more efficient that re-running the conventional (offline) analysis every time new sequences are added. Improving the method’s efficiency further is an interesting future direction. Our method also extends to a wide range of evolutionary models – all those available in BEAST 2 and based on “single tree” models, that is, excluding e.g. the multi-species coalescent based models (Douglas et al., 2022). Extending the method to further evolutionary models such as the multi-species coalescent is also an interesting future direction.

Motivated by a number of successes of MCMC-based phylogenetics in the online framework, including the one presented here, we finish this paper by discussing online approaches to other phylogenetic methodologies. Specifically, we first suggest a simple way to extend our results to the Maximum Likelihood paradigm in phylogenetics, and second we argue that designing online distance-based phylogenetic inference approaches might require significant theoretical advances in algorithms and data structures.

### 5.1. Online maximum likelihood tree reconstruction methods

Maximum likeli-hood methods and maximum *a posteriori* (MAP) algorithms can benefit from the same approach as the Bayesian online approach presented here by using the first two phases of our algorithm – adding taxa followed by local tree optimisation – and taking this as a starting point to continue optimisation using standard methods.

This is implemented in BEAST 2 (BEASTLabs package^2^), for example, using the simulated annealing approach (Van Laarhoven and Aarts, 1987) using MCMC proposals for suggesting new states. After running simulated annealing on a set of taxa, a state file with the best state found so far is produced. The `StateExpander` application in the online package^3^ can then be used to expand this state with new sequences by running the first two phases of the Bayesian online algorithm, but only on the saved state, not on a set of states like in the Bayesian online algorithm. Simulated annealing can then be resumed with the expanded state as a starting point, incorporating new sequences.

A tutorial for using online simulated annealing implemented in BEAST 2 is available at https://github.com/rbouckaert/online-tutorial.

### 5.2. Alternative use cases

Here, we describe a number of practical uses of the individual steps of our online MCMC algorithm, in addition to the conventional use case presented in previous sections.

#### Altering models

It is not uncommon to run a Bayesian analysis on a single alignment with different substitution models, clock models, or tree priors to establish model (in)sensitivity. This is a scenario where the data remains the same, but parameters of models may need to be added or removed, so it makes little sense to run the first and second phase of the algorithm. However, running the third phase of the algorithm can be done in parallel and in addition to starting with a good tree and other shared parameters can benefit from auto-optimised MCMC operator settings obtained from previous runs and paralellism, thus substantially reducing computation time.

#### MCMC starting state

If one does not have sufficient computational resources to run many threads in parallel, simple MCMC will have to suffice. However, one can expect trees and joint parameters to be not too different between different analyses with different models, and it is still possible to reuse the end state and operator settings obtained from one analysis as starting point for the next analysis. When the set of taxa changes, running the first two phases of Algorithm 1 still makes sense. The `StateExpander` utility in the online package for BEAST 2 can be used to produce a good starting state this way.

#### Resuming

.Since human inspection of traces appears to be the most reliable way to determine convergence of an MCMC analysis at the moment, the algorithm can stop after a fixed number of MCMC steps have been performed. This may not be sufficient to reach convergence, and the parallel chains can then be resumed at that stage by running `TraceExpander`^4^.

#### Breaking up large analyses

When an analysis with a large number of taxa is performed, it can take a long time, sometimes weeks, for plain MCMC to converge. In the following, we assume computational time of an MCMC analysis tends to be roughly cubic in the number of taxa. By randomly subsampling the taxa, first a smaller analysis can be performed on the random subset, which should take significantly less time: taking a quarter of the taxa would result in an analysis taking around 1*/*4^3^ = 1*/*64 of the time of a full analysis. New taxa can be added using our online algorithm, which due to parallelisation requires less computational time: adding another quarter of the taxa running on 10 threads requires 1*/*2^3^ ∗ 1*/*10 = 1*/*80 of a full run after burn-in. The process can be repeated until all taxa are included in the analysis. Total run time when splitting in 4 and running on 10 threads would be approximately 1*/*64+1*/*80+27*/*80+1*/*10 = 0.46 times the run time of a full run. A further benefit is that analyses on subsets of taxa will be available earlier in the process, which can give preliminary insights into the full analysis, and allow earlier refinements in, for example, model selection.

For convergence analysis, it is good practice when running an MCMC analysis to run multiple instances in order to be sure that each of them converge to the same distribution. Likewise, we recommend to run at least three, preferably more, instances of the online algorithm in parallel to allow judging how much the distributions agree. If there is significant deviation, the chain either has not run for long enough, or the analysis just does not converge to the correct distribution due to multi-modality caused by, for instance, overparameterisation.

A tutorial for these alternative use cases implemented in BEAST 2 is available at https://github.com/rbouckaert/online-tutorial.

### 5.3. Online distance-based tree reconstruction methods

A natural question that arises in the context of presented online approaches to Bayesian and Maximum Likelihood phylogenetic inference is whether or not distance-based tree reconstruction methods have efficient online versions. Here, we briefly consider two such methods, namely the Unweighted Pair Group Method with Arithmetic Means (UPGMA) (Sokal, 1958) and Neighbour Joining (NJ) (Saitou and Nei, 1987).

The art of designing online algorithms often relies on identifying a distance measure between offline algorithm’s outputs under which the algorithm is “stable” (Gavryushkin et al., 2015, 2016). Extended to distance-based phylogenetics, this task requires identifying a neighbourhood of the running tree that restricts the range of possible trees the running tree can attain following an update operation. In our context, this is equivalent to the following. Given a UPGMA (NJ) tree *T* reconstructed from a set of sequences (taxa) *S*, find the range of possible UPGMA (NJ) trees *T*′ reconstructed from a set of sequences *S* ∪{*x*}. Figure 5 summarises our computational assessment of this range.

**Figure 5.**
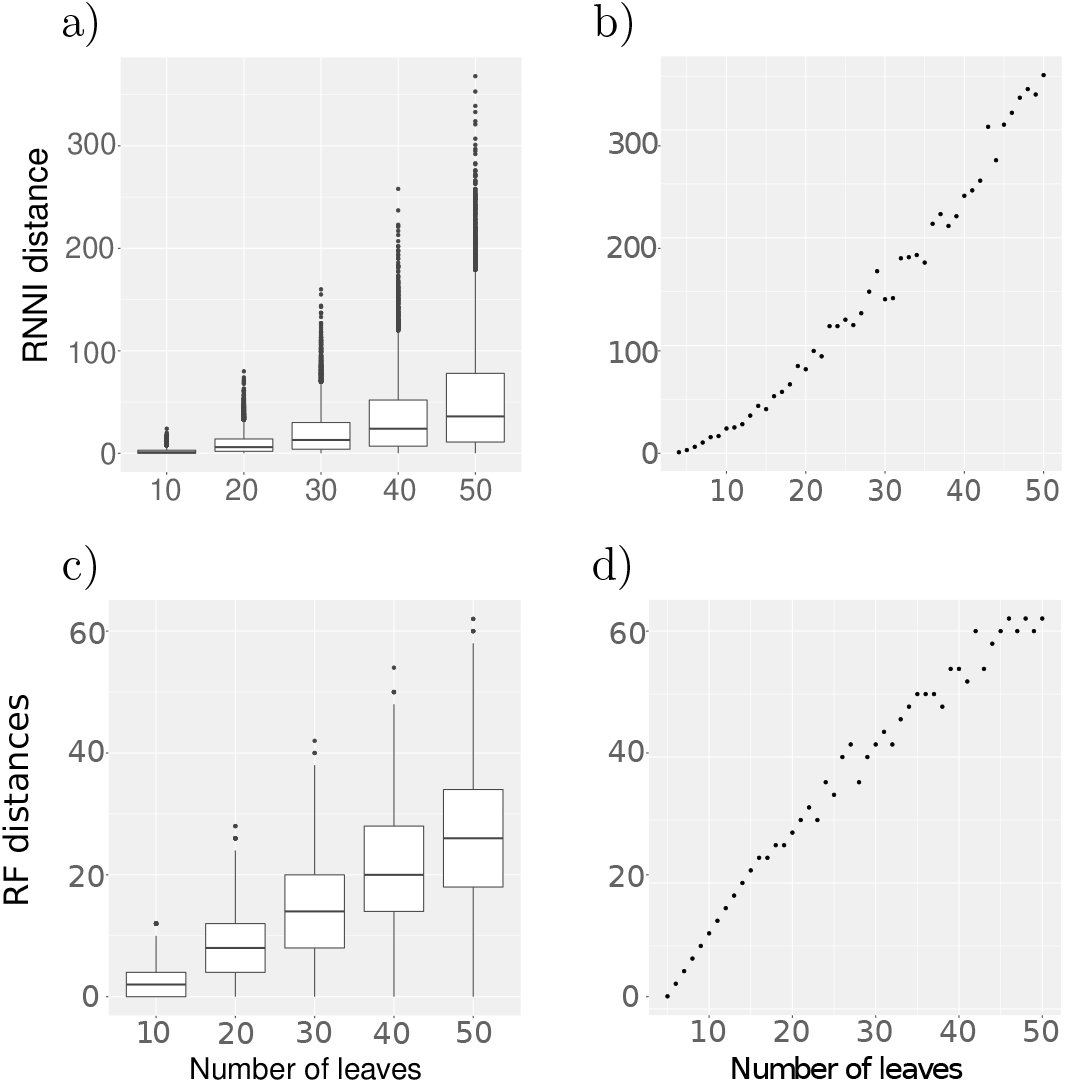
RNNI distances between UPGMA trees (top) and Robinson Foulds distances between NJ trees (bottom) *T* and *T*′ for *n ∈ {* 10, 20, 30, 40, 50} on the left. Shown are the distances between *n* simulated points in the Euclidean space, with 10, 000 tree pairs for each value of *n*. On the right the maximum RNNI and RF distances for the same simulations with *n* = 4, …, 50 are plotted.

The results presented in Figure 5 are obtained by simulating a set *S ∪ {x}* of random points in the 50-dimensional Euclidean space [0, 1]^50^. The matrix of their pairwise distances is then used as input to the UPGMA (NJ) algorithm, resulting in a tree *T* from which a random taxon *x* is deleted. This tree is then compared with the tree *T*′ computed by UPGMA (NJ) for the matrix of pairwise distances of points in *S*. For this comparison we use the RNNI distance (Gavryushkin et al., 2018; Collienne and Gavryushkin, 2021; Collienne, 2021) for UPGMA trees and the Robinson-Foulds distance ((Robinson and Foulds, 1981)) for NJ trees (see Supplementary Material for details).

As these results suggest, there is no obvious restriction on the range possible trees can attain, meaning the algorithm is likely to be not stable, so an online version of the algorithm would be hard to design. Specifically, an efficient online version of the UPGMA (NJ) algorithm for the general case of arbitrary distance matrices would either require an entirely novel distance measure between trees under which the changes become local or an entirely novel approach to online algorithms not based on the idea of stability. Both of these tasks appear to be hard.

One alternative formulation of an online distance-based phylogenetic reconstruction method could be to consider distance matrices corresponding to sequence alignments, rather than arbitrary Euclidean matrices as we considered above. In Figure 6 we repeated the same computational experiment as above but generated the distance matrices by simulating sequences of length 1, 000 for a random Yule-Harding tree on *n* leaves, assuming an HKY model (Hasegawa et al., 1985) with transition/transversion ratio of 2 and base frequencies 0.1, 0.2, 0.3, 0.4 for A, C, G, T, respectively. We then used this alignment to produce a distance matrix *D* and followed the approach above. Further details on this simulation are provided in Supplementary Material.

**Figure 6.**
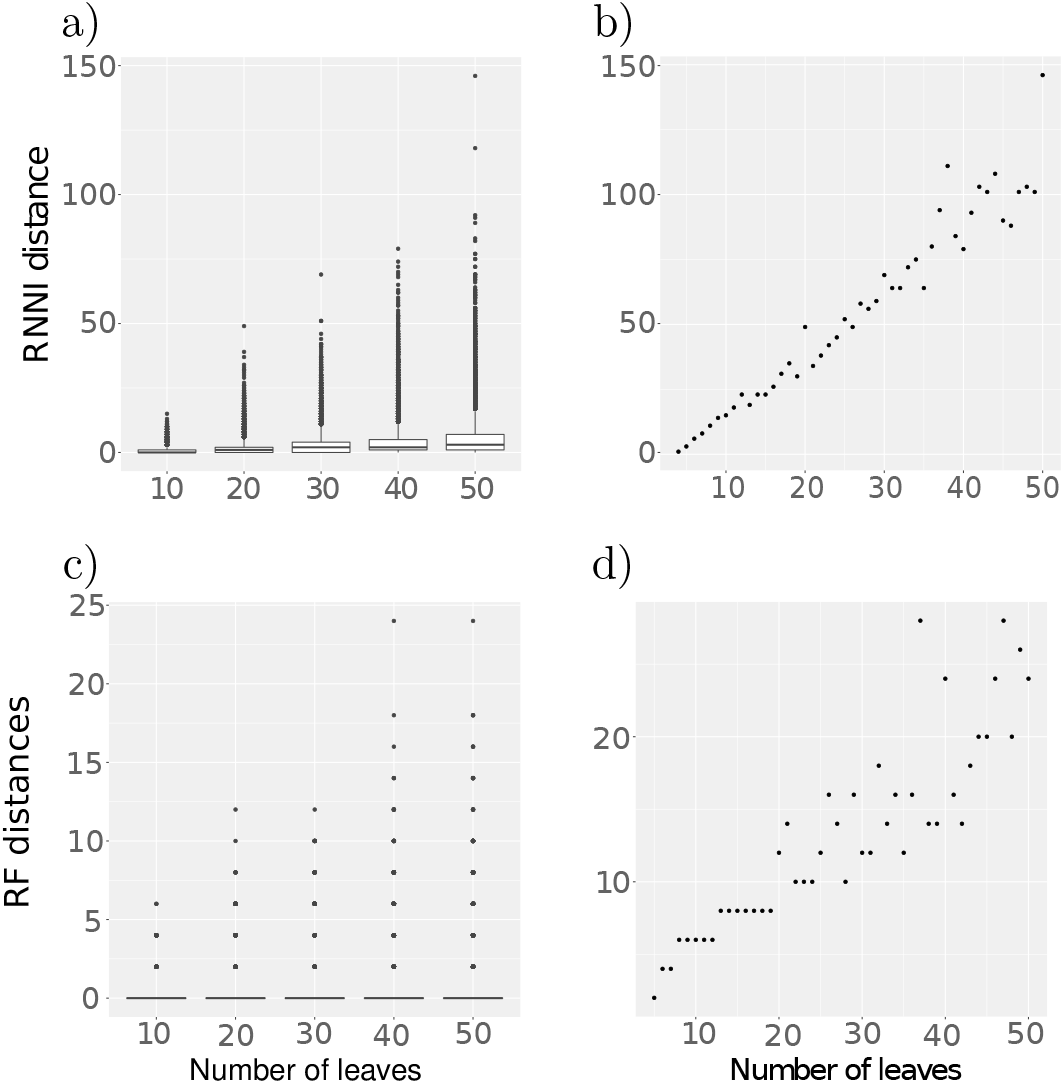
RNNI distances between UPGMA trees (top) and Robinson Foulds distances between NJ trees (bottom) *T* and *T*′ for *n ∈ {* 10, 20, 30, 40, 50} on the left. Shown are the distances between *n* simulated DNA sequences, with 10, 000 tree pairs for each value of *n*. On the right the maximum RNNI and RF distances for the same simulations with *n* = 4, …, 50 are plotted.

This result suggests that restricted to the class of distance matrices corresponding to sequence alignments, there might be a way to formulate a local condition to isolate a small enough range of trees that can be attained by the UPGMA (NJ) algorithm and leverage this stability property to design an efficient online algorithm. A potentially non-trivial task for this approach, however, would be to ensure the condition of the distance matrix remaining “alignment-like” is satisfied, and understanding the behaviour of the algorithm when this condition is violated.

## 6 Availability

The open source `online` package for BEAST 2 (Bouckaert et al., 2019) is available under GPL at https://github.com/rbouckaert/online. It supports adding and deleting taxa, as well as changing models. The analysis can be resumed using full MCMC only if at first it turns out that the algorithm has not converged. A tutorial for the package is available at https://github.com/rbouckaert/online-tutorial.

## S1. Supplementary Material

### S1.1. Stability analysis for distance-based methods

Here we present details on the simulation study analysing the stability of distance-based tree inference methods (Section 5.3). We investigate the stability of two distance-based tree inference methods, namely Unweighted Pair Group Method with Arithmetic Means (UPGMA) (Sokal, 1958) and Neighbour Joining (NJ) (Saitou and Nei, 1987). These methods take distance matrices, which are usually derived from sequence alignments, as input and produce a phylogenetic tree as output. Our simulations are implemented mostly in R, using the packages *phangorn* (Schliep, 2011) *and ape* (Paradis and Schliep, 2019), and for simulating alignments we use the tool *AliSim* from the software package *IQ-TREE* (Ly-Trong *et al*., 2021). All implementations can be found in (Collienne, 2022). We simulate distance matrices in two different ways (Sections S1.1.1 and S1.1.2) to assess the stability of distance based tree inference methods.

#### S1.1.1. Euclidean distance matrices

In our first approach we simulate *n* points in the 50-dimensional space [0, 1]^50^ and compute a Euclidean distance matrix *D* for those *n* points, which represent *n* taxa. We then delete one column and row of *D* corresponding to a randomly chosen taxon *x*, giving us a matrix *D* ↾ *n −* 1 for *n −* 1 taxa. The two distance matrices are then used to compute trees *Tree*(*D*) and *Tree*(*D* ↾ *n −* 1) (using UPGMA (*phangorn*) and NJ (*ape*)). To analyse the influence of deleting a taxon of the reconstructed tree, and hence the stability of the two algorithms, we compare *Tree*(*D* ↾ *n −* 1) with *Tree*(*D*) ↾ *n −* 1, which results from deleting the taxon from *Tree*(*D*). For this comparison we use the RNNI distance (Collienne and Gavryushkin, 2021; Collienne, 2021) for UPGMA trees and the Robinson-Foulds distance (as implemented in *phangorn*) for NJ trees. We have reverted to the Robinson-Foulds distance for unrooted trees because no tree rearrangement-based distance is practically computable for such trees. The experiment is repeated 10, 000 times for trees on *n* = 50 leaves for both NJ and UPGMA. The results are presented in Figure S1, where the distributions of distances normalised by the corresponding space’s diameters are shown.

The results in Figure S1 suggest that the distance between trees *Tree*(*D* I *n −* 1) and *Tree*(*D*) ↾ *n −* 1 cannot be bounded by a constant. This indicates that neither UPGMA nor NJ are stable under the distance measures RNNI and Robinson-Foulds.

We continue our stability analysis by varying the number of taxa *n* and repeating the simulation as described above. On the left of Figure 5 we show the RNNI distances between UPGMA trees (top) and Robinson-Foulds distances between NJ trees (bottom) for 10, 000 repetitions of our simulations for each value of *n ∈ {*10, 20, 30, 40, 50}. These plots reveal that with increasing *n* the RNNI distance between the trees *Tree*(*D*) I *n* 1 and *Tree*(*D* ↾ *n −* 1) is unbounded, too. This can also be seen in the plots on the right of Figure 5, where the maximum RNNI and RF distances for 10, 000 repetitions of the simulation (as described above) for each *n* = 4, …, 50 are shown. This in particular implies that there is no constant bound on the distance between these trees.

#### S1.1.2. Distance matrices from simulated alignments

In practice, distance matrices used to infer phylogenetic trees are generated from sequence alignment, so they form a subclass of the class of all distance matrices. Therefore, a possible way forward to design an online UPGMA (or NJ) algorithm can be to assume that the input distance matrix is of that sort. We now hence continue our stability analysis by considering distance matrices corresponding to sequence alignments.

**Figure S1.**
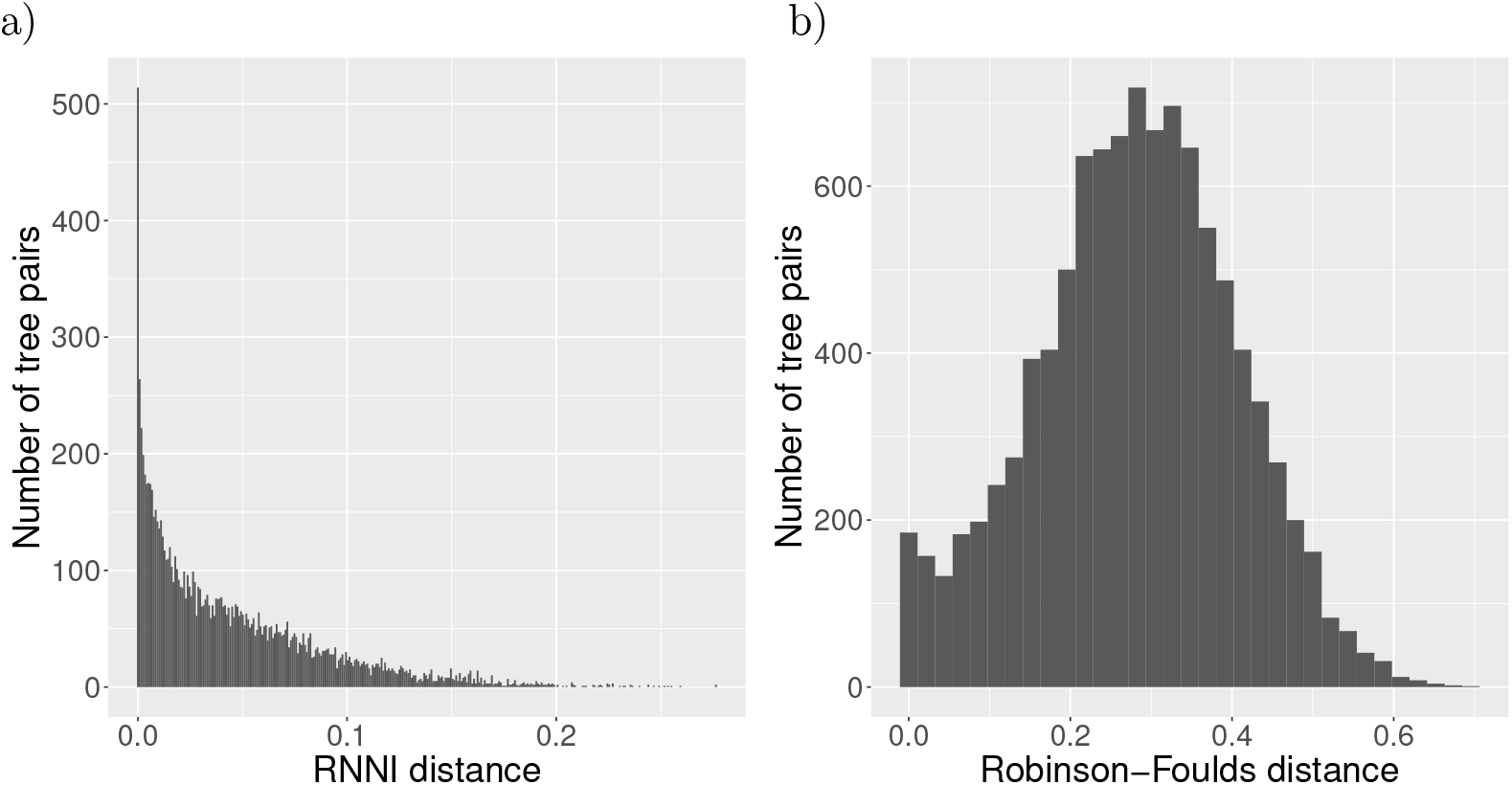
Distances between trees *Tree*(*D*) ↾ *n −* 1 and *Tree*(*D* ↾ *n −* 1) with *n* = 50 leaves where *D* represents distances between points in a cube. Normalised (by the diameter of the space) RNNI distance between UPGMA trees on the left and normalised (by the diameter of the space) Robinson-Foulds distance between NJ trees on the right. Shown is the number of tree pairs out of a total of 10, 000 simulations (*y*-axis) with distance *d* (*x*-axis).

For simulating sequence alignments we use the tool *AliSim* from the software package *IQ-TREE* (Ly-Trong et al., 2021). *Sequences of length 1*, 000 are computed for a random Yule-Harding tree on *n* leaves, assuming an HKY model (Hasegawa et al., 1985) with transition/transversion ratio of 2 and base frequencies 0.1, 0.2, 0.3, 0.4 for A, C, G, and T, respectively. This results in an alignment *A* of *n* DNA sequences. From this alignment we generate a second alignment *A* ↾ *n −* 1 of *n −* 1 DNA sequences, which results from randomly choosing a taxon *x* and deleting its sequence from *A*. In the next step we compute distance matrices *D* and *D* ↾ *n* 1 for both sequence alignments with the function dist.ml (assuming an F81 model (Felsenstein, 1981)) from the R package *phangorn*. For those distance matrices we then compute UPGMA and NJ trees *Tree*(*D* ↾ *n −* 1) and *Tree*(*D*) ↾ *n −* 1 as we did above.

In Figure S2 one can see the RNNI and Robinson-Foulds distances between trees *Tree*(*D* ↾ *n −* 1) and *Tree*(*D*) ↾ *n −* 1 from our second simulation using sequence alignments. One can see that the distances in this simulations are in general smaller than in the ones presented in Figure S1.

For this simulation of distance matrices from alignments, we also compared RNNI distances (for UPGMA) and RF distances (for NJ) of trees for a varying number of taxa. We therefore repeat the simulations of distance matrices from alignments in the same way as before, but repeat it for a varying number of taxa. On the left of Figure 6 we show the RNNI distances between UPGMA trees (top) and Robinson-Foulds distances between NJ trees (bottom) for 10, 000 repetitions of our simulations for each value of *n ∈ {*10, 20, 30, 40, 50}. These plots reveal that with increasing *n* the RNNI distance between the trees *Tree*(*D*) ↾ *n −* 1 and *Tree*(*D* ↾ *n −* 1) is unbounded, too. Again, the distances are smaller than the ones in Figure 5, but they are still unbounded. This can also be seen in the plots on the right of Figure 5, where the maximum RNNI and RF distances for 10, 000 repetitions of the simulation (as described above) for each *n* = 4, …, 50 are shown. The distances between the trees in this second simulations where distances come from simulated alignments are in general smaller than in our first simulation in Section S1.1.1, but they are unbounded as well. This suggests that there is no constant bound on the distance between these trees.

**Figure S2.**
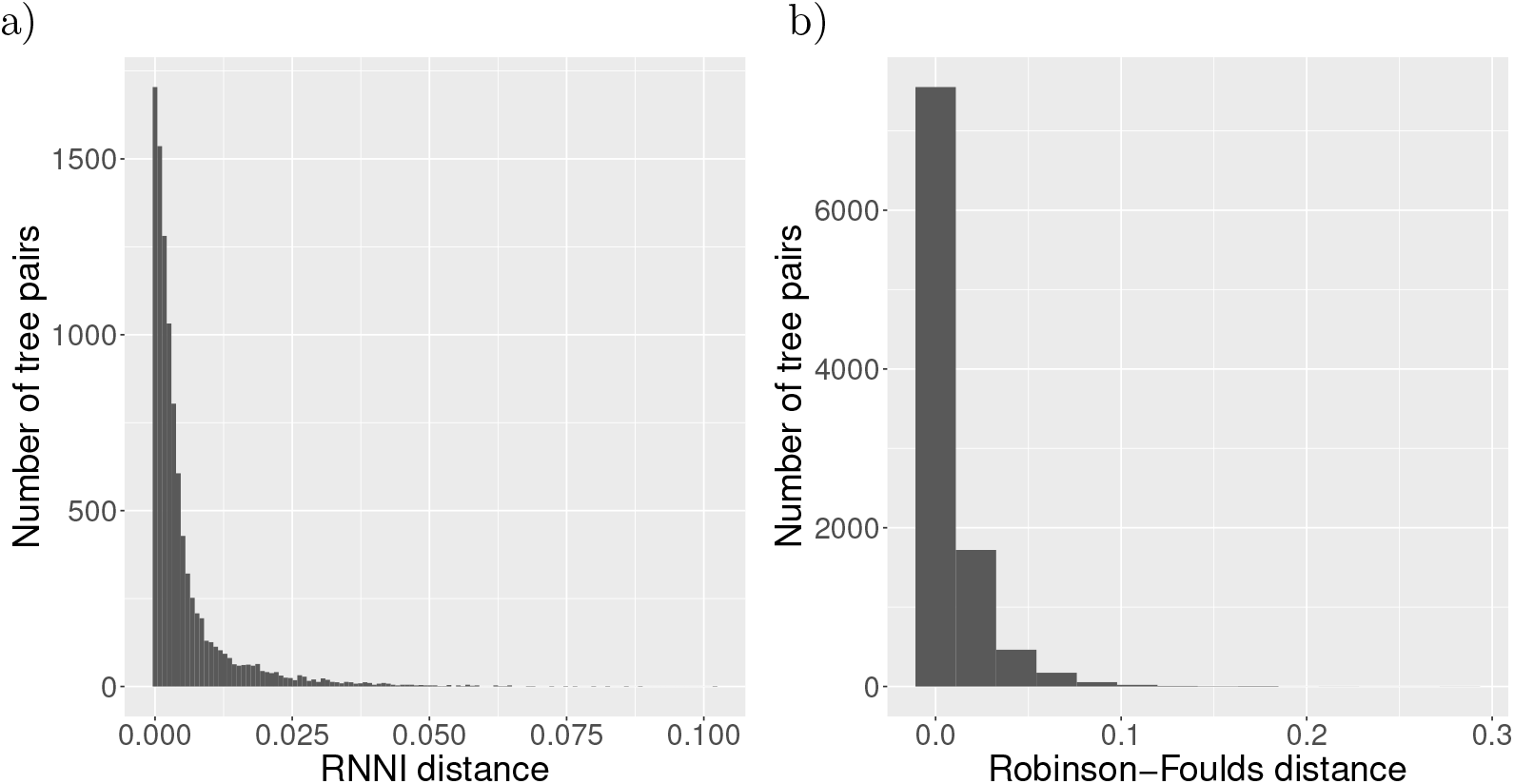
Distances between trees *Tree*(*D*) ↾ *n −* 1 and *Tree*(*D* ↾ *n −* 1) on *n* = 50 where *D* represents distances between sequences of an alignment. Normalised RNNI distance between UPGMA trees on the left and normalised Robinson-Foulds distance between NJ trees on the right. Shown is the number of tree pairs out of a total of 10, 000 simulations (*y*-axis) with distance *d* (*x*-axis).

## S2. Supplementary figure

**Figure S3.**
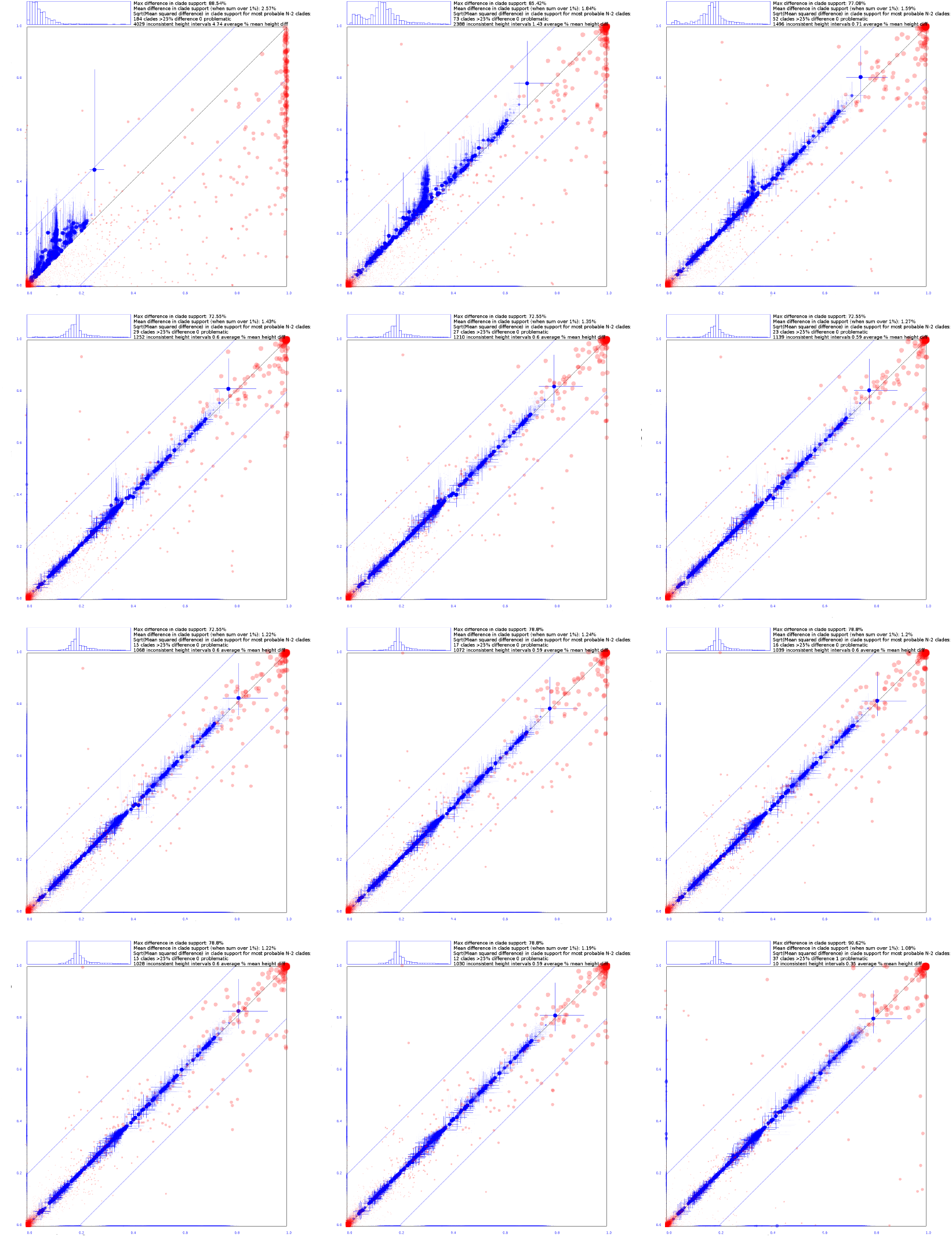
As Fig. 4 but including comparison after 1 up to 11 online iterations (in reading order) on the vertical axis. The last panel compares two independent MCMC runs for reference.

## Footnotes

1 Sourced from http://wiki.christophchamp.com/index.php/NEXUS_file_format

2 https://github.com/BEAST2-Dev/BEASTLabs/

3 https://github.com/rbouckaert/online

4 Without specifying xml2.

## Notes

AG and LC acknowledge support from the Royal Society Te Apārangi through a Rutherford Discovery Fellowship (RDF-UOO1702). This work was partially supported by Ministry of Business, Innovation, and Employment of New Zealand through an Endeavour Smart Ideas grant (UOOX1912) and a Data Science Programmes grant (UOAX1932).

https://github.com/rbouckaert/online

### Competing Interest Statement

The authors have declared no competing interest.

